# Placental microRNA signatures of spontaneous preterm birth

**DOI:** 10.64898/2026.08.21.746278

**Authors:** Mariana Parenti, Elizabeth M. Kennedy, Evan J. Firsick, Samantha Lapehn, James MacDonald, Theo Bammler, Daniel A. Enquobahrie, Kaja Z. LeWinn, Nicole R. Bush, Stephen A. McCartney, Carmen J. Marsit, Qi Zhao, Sheela Sathyanarayana, Alison G. Paquette

## Abstract

**Background:** The placenta has a unique transcriptomic profile, including microRNAs that are secreted into maternal circulation throughout pregnancy. MicroRNAs are small, non-coding RNA that post-transcriptionally regulate gene expression. Spontaneous preterm birth (sPTB) is associated with substantial differences in both placental pathophysiology and placental gene expression compared to term birth. We aimed to generate microRNA signatures of sPTB and map them to target genes using a microRNA–mRNA network.

**Methods:** This study was conducted within the Conditions Affecting Neurocognitive Development and Learning in Early childhood (CANDLE) study. Placental samples were collected at delivery, and RNA was isolated for mRNA and microRNA sequencing. To investigate sPTB, this study excluded placental samples of participants with iatrogenic indications for PTB or induced labor. We examined differences in microRNA expression in participants who delivered before 37 weeks (*N*=35) compared to term participants (*N*=404) in a series of covariate-adjusted linear regression models. We used paired placental microRNA and mRNA expression data from this cohort to validate associations between computationally predicted microRNA–mRNA pairs and establish a microRNA–mRNA network.

**Results:** Expression of 7 microRNAs were increased in sPTB (FDR<0.05) and were inversely correlated with sPTB-associated genes involved in immune signaling. Expression of 12 microRNAs were decreased in sPTB, including 4 members of the maternally expressed chromosome 14 microRNA cluster (miR-376a-3p, miR-376c-3p, miR-377-3p, and miR-381-3p). These microRNAs were predicted to negatively regulate oxidative phosphorylation genes that were increased in sPTB. The associations between miR-376c-3p and miR-377-3p and oxidative phosphorylation were confirmed in microRNA knockdown experiments.

**Conclusions:** This study highlights potential biological mechanisms by which placental microRNA dysfunction might contribute to sPTB and highlights putative sPTB biomarkers that may be detectable in maternal circulation.

## 1. INTRODUCTION

Preterm birth (**PTB**, birth before 37 weeks’ gestation) affects 1 in 10 infants born in the United States and is a leading cause of infant morbidity and mortality [1,2]. While some PTBs are medically indicated for conditions like preeclampsia or fetal growth restriction, approximately 60% of PTBs occur spontaneously due to preterm premature rupture of membranes or preterm labor [3]. Even small increases in gestational length are associated with improved survival and health outcomes in both preterm and early-term (between 37–39 weeks’ gestation) infants compared to full-term (39–41 weeks’ gestation) infants [4–7]. Thus, it is critically important to understand the underlying mechanisms contributing to spontaneous PTB (**sPTB**), as well as variability in gestational length. sPTB is a syndrome with multifactorial causes that converge in maternal immune activation, inflammation, and oxidative stress, leading ultimately to premature delivery [8].

As the site of the maternal–fetal interface, the placenta is uniquely situated to coordinate the molecular mechanisms underlying sPTB. The placenta influences maternal metabolism, modulates immune responses, facilitates maternal-fetal crosstalk, and serves as the source of nutrition for the developing fetus [9]. Regulation of placental biology must be dynamic and finely tuned to support fetal development and appropriately timed birth. Thus, signatures of regulatory mechanisms influencing placental function are a promising source of insight into sPTB etiology.

MicroRNAs (**miRNAs**) are a class of endogenous, small non-coding RNAs that post-transcriptionally regulate gene expression [10]. These single-stranded miRNAs bind to target gene transcripts, which inhibits translation, promotes mRNA degradation, and usually silences the target gene [10]. miRNAs play critical roles in regulating development and cell differentiation, starting in the pre-implantation embryo [11]. Across gestation, miRNAs exhibit dynamic patterns of expression as the placenta responds to changes in oxygen tension and fetal growth [12,13]. miRNAs regulate core aspects of trophoblast function, including invasion, migration, proliferation, and differentiation into extravillous trophoblasts and syncytiotrophoblasts, as well as cellular metabolism and placental angiogenesis (reviewed in [14]).

Notably, a number of miRNAs are predominantly or exclusively expressed in the placenta, including the Chromosome 14 miRNA cluster (**C14MC**) and Chromosome 19 miRNA cluster (**C19MC**). Both clusters are imprinted, with the C19MC miRNAs expressed only from the paternal allele and the C14MC miRNAs expressed only from the maternal allele [15,16]. The C14MC is eutherian-specific, with highly conserved orthologs across placental mammals [17]. The evolutionarily conserved target genes of the C14MC are enriched in biological processes related to mRNA processing, embryonic development, brain development, and axon guidance [17]. This cluster is also expressed in the postnatal brain, where the miRNAs regulate genes involved in transcriptional and developmental regulation [18]. C19MC is primate-specific [16], and C19MC miRNAs might contribute to maintaining maternal-fetal tolerance by interfering with the innate immune response in trophoblasts [19]. Both clusters are closely related to cell fate in the early embryo, with C19MC highly expressed in the trophectoderm and C14MC highly expressed in the inner cell mass [11]. As gestation advances, C14MC miRNAs are expressed in the fetus and placenta and enter into maternal circulation, while C19MC miRNAs are expressed in the placenta and enter into both the fetal and maternal compartments [20,21]. As a result, pregnancy-specific miRNAs could serve as biomarkers of placental function that can be monitored in maternal circulation [22].

miRNA signatures have been profiled in various tissues to serve as biomarkers and illuminate the mechanisms of sPTB. We have previously identified miRNA signatures of spontaneous preterm labor in maternal whole blood and monocytes, including C14MC miRNAs [23]. Other studies have used circulating miRNAs in blood collected before 20 weeks’ gestation to predict sPTB [24–30]. Despite the importance of predictive biomarkers, a functional understanding of how miRNA regulation might contribute to placental mechanisms of sPTB is still needed. Some studies have identified differentially expressed miRNAs in placental tissue and fetal membranes [31–34]. However, most analyses have used targeted PCR-based or array-based approaches to measure miRNAs without paired mRNA data, limiting their ability to investigate putative regulatory mechanisms. This is especially important in the placenta, given its unique miRNA repertoire. Paired placental miRNA and mRNA signatures have been used to investigate risk factors associated with PTB in a cohort of extremely preterm infants [35]. We have previously established a transcriptomic signature of sPTB to further our understanding of the underlying placental mechanisms of sPTB [36]. In that study, sPTB was associated with decreased placental gene expression in signaling pathways related to placental function and growth [36].

The goal of this study was to identify placental miRNA signatures of sPTB and link them to their sPTB-associated target genes in a large, richly characterized pregnancy cohort. We used unbiased genome scale RNA sequencing to quantify placental expression of over 500 miRNAs and identify those that were differentially expressed in sPTB compared to placentas delivered at term. We used the paired expression data from the same participants to generate a miRNA–mRNA network to better characterize the regulatory functions of placental miRNAs associated with sPTB and gestational length. We hypothesized that the placental miRNA signatures of sPTB would include placenta-specific miRNA species and would regulate genes associated with sPTB.

## 2. METHODS

All research activities for the Conditions Affecting Neurocognitive Development and Learning in Early Childhood (**CANDLE**) Study were approved by the University of Tennessee Health Sciences Center IRB, the ECHO prenatal and early childhood pathways to health consortium (**ECHO-PATHWAYS**) single IRB [37,38], and Seattle Children’s Research Institute (STUDY00004005).

### 2.1. Participants and data collection

This analysis was conducted using samples collected as part of the CANDLE study, a prospective pregnancy cohort conducted in Shelby County, Tennessee between December 2006 and July 2011 [37]. Pregnant participants were recruited during their second trimester and were considered eligible if they were between 16 and 28 weeks of gestation, had an uncomplicated singleton pregnancy, and planned to give birth at a participating Shelby County health care center [37]. Participants with placental RNA sequencing and small non-coding RNA sequencing data were included in this analysis. In this analysis, we excluded samples from pregnancies with medical indications for preterm delivery <37 week’s gestation, including placental abruption, preeclampsia, hypertension, or chorioamnionitis (**Figure S1**). We used paired placental RNA sequencing and small non-coding RNA sequencing data from this subset of the CANDLE cohort to validate computationally predicted mRNA targets of placental miRNAs. To identify miRNAs whose placental expression was associated with sPTB, we further excluded participants with induced labor. The final analytic population included 35 sPTB and 404 term placental samples. Covariate data were collected by maternal self-report or medical record abstraction [37]. Smoking status was defined as either prenatal urinary cotinine >200 ng/mL [39] or self-reported smoking during pregnancy.

### 2.2. Placental sample collection

As previously described, placental tissue was collected by CANDLE researchers within 15 minutes of delivery. A piece of placental villous tissue approximately 2 cm × 0.5 cm × 0.5 cm was dissected from the middle of the placental parenchyma [36,40,41]. The tissue was further split into four cubes, which were refrigerated in RNALater at 4°C overnight, transferred to fresh RNALater, and stored at -80°C. The tissue was manually dissected to remove maternal decidual tissue and fetal villous tissue was used for RNA isolation as previously described [40]. Briefly, approximately 30 mg of tissue was homogenized using a TissueLyser LT instrument (Qiagen) and RNA and miRNAs were isolated using the AllPrep DNA/RNA/miRNA Universal Kit (Qiagen). Only samples with RNA integrity number >7 as measured using a Bioanalyzer 2100 with RNA 6000 Nanochips (Agilent) were sequenced.

### 2.3. mRNA sequencing

The ECHO-PATHWAYS consortium generated placental RNA sequencing data from 794 participants in the CANDLE cohort as described by LeWinn et al [38]. RNA sequencing was conducted at the University of Washington Northwest Genomics Center as previously described [36,40,41]. Total RNA was poly-A enriched, complementary DNA libraries were prepared using the TruSeq Stranded mRNA kit (Illumina), and each library was sequenced to an approximate depth of 30 million reads on an Illumina HiSeq 4000 instrument. RNA sequencing data quality control was performed using both the FASTX-tool (version 0.0.13) and FastQC (version 0.11.2) toolkits [42]. Transcript abundances were estimated by aligning to the GRCh38 transcriptome (Gencode version 33) using Kallisto [43], then collapsed to the gene level using the Bioconductor tximport package [44] and scaled to the average transcript length.

### 2.4. Small non-coding RNA sequencing

Small non-coding RNA (sncRNA) sequencing was conducted using total RNA from 765 samples that were previously subjected to mRNA sequencing. A total of 100 ng total RNA was used for miRNA sequencing library preparation with the QIAseq miRNA Library Kit (Qiagen). Adaptors were ligated first to the 3’ end of the sncRNAs, followed by adaptor ligation at the 5’ end. In a reverse transcription step, unique molecular indexes (**UMIs**) were introduced, and the ligated miRNAs were converted to cDNA. Subsequently, cDNA cleanup, library amplification, and library cleanup were performed. The libraries were sequenced on an Illumina NextSeq 2000 instrument with a single read length of 75 bp and dual index reads of 10 bp. Raw data were de-multiplexed and FASTQ files for each sample were generated using Illumina bcl2fastq (version 2.20.0.422). FASTQ files were uploaded to GeneGlobe Data Analysis Center (Qiagen), which was used for quality control, alignment to a miRNA reference (miRbase, version 22, *Homo sapiens*), and collapsing UMI reads to counts.

### 2.5. Statistical Analyses

We compared the population of spontaneous term and preterm births (*N*=439) included in this analysis to the remainder of the CANDLE cohort (**Table S1**), as well as the spontaneous term births to sPTB (**Table 1**). Differences in population characteristics were analyzed using *χ*^2^ tests for categorical variables and two-sample T tests for continuous variables. All analyses were conducted using the *R* statistical language (R Core Team, Vienna, Austria).

**Table 1.** Participant characteristics by sPTB birth outcome.

|  | **Term (N=404)** | **Preterm (N=35)** | **P value** |
| --- | --- | --- | --- |
| **Maternal age at birth** |  |  | 0.099 |
| Mean (SD) | 27.9 (5.2) | 26.4 (5.7) |  |
| **Maternal pre-pregnancy BMI** |  |  | 0.053 |
| Mean (SD) | 27.6 (7.4) | 25.1 (6.2) |  |
| **Maternal education** |  |  | 0.453 |
| Less than high school | 26 (6.4%) | 5 (14.3%) |  |
| High school graduate/GED | 175 (43.3%) | 16 (45.7%) |  |
| Technical school | 35 (8.7%) | <5 |  |
| College degree | 107 (26.5%) | 7 (20.0%) |  |
| Grad/Professional degree | 61 (15.1%) | <5 |  |
| **Estimated total annual household income at enrollment** |  |  | 0.042 |
| \$0-4999 | 44 (10.9%) | 5 (14.3%) | |
| \$5000-9999 | 29 (7.2%) | 7 (20.0%) | |
| \$10000-14999 | 23 (5.7%) | <5 | |
| \$15000-19999 | 24 (5.9%) | <5 | |
| \$20000-24999 | 18 (4.5%) | 5 (14.3%) | |
| \$25000-34999 | 46 (11.4%) | <5 | |
| \$35000-44999 | 33 (8.2%) | <5 | |
| \$45000-54999 | 45 (11.1%) | <5 | |
| \$55000-64999 | 19 (4.7%) | <5 | |
| \$65000-74999 | 40 (9.9%) | <5 | |
| \$75000 or over | 83 (20.5%) | 6 (17.1%) | |
| **Maternal self-reported race** |  |  | 0.309 |
| Black/African American | 214 (53.0%) | 23 (65.7%) |  |
| White | 167 (41.3%) | 9 (25.7%) |  |
| Asian | 5 (1.2%) | <5 |  |
| Other | <5 | <5 |  |
| Multiple race | 17 (4.2%) | <5 |  |
| **Maternal smoking status <sup>a</sup>** |  |  | 0.701 |
| Never | 361 (88.6%) | 32 (91.4%) |  |
| Previous or current smoker | 43 (10%) | <5 |  |
| **Maternal self-reported alcohol use during pregnancy** |  |  | 0.302 |
| No | 358 (88.6%) | 33 (94.3%) |  |
| Yes | 46 (11.4%) | <5 |  |
| **Fetal sex** |  |  | 0.607 |
| Male | 201 (49.8%) | 19 (54.3%) |  |
| Female | 203 (50.2%) | 16 (45.7%) |  |
| **Labor type** |  |  | 0.004 |
| Spontaneous | 114 (28.2%) | 13 (37.1%) |  |
| Spontaneous, augmented | 174 (43.1%) | 21 (60.0%) |  |
| No labor | 116 (28.7%) | <5 |  |
| **Delivery method** |  |  | 0.008 |
| Vaginal | 231 (57.2%) | 28 (80.0%) |  |
| C-section | 173 (42.8%) | 7 (20.0%) |  |

Continuous variables are reported as mean and standard deviation (SD) and differences were evaluated using two-sample T tests. Categorical variables are reported as N (%) and were tested using $\chi^2$ tests.
<sup>a</sup> Maternal smoking status was composite variable determined by self-report and urinary cotinine where available. Participants who self-reported ever smoking or who had urinary cotinine > 200 ng/mL were classified as previously or currently smokers.

### 2.6. Placental miRNA–mRNA network construction

We constructed a miRNA–mRNA network using computationally predicted miRNA–target gene pairs and validated these associations in the placenta using paired sequencing data from the CANDLE cohort. We used 2 databases of computationally predicted miRNA–target gene interactions (TargetScan Human and miRDB) to identify putative miRNA–mRNA pairs [45,46]. We leveraged all participants with paired miRNA and mRNA expression data (*N*=691) to confirm these pairs based on co-expression and establish the network (**Figure S1**). We filtered mRNA sequencing data to only include protein-coding genes and removed mRNA with low expression (mean log counts per million (logCPM<0). Scaling factors were calculated using the trimmed mean of M-values (TMM) as previously described [36,40,41]. Small RNA UMI count data were similarly filtered to retain only miRNA, filtered to remove logCPM<0, and TMM scaling factors were calculated. Batch effects were removed from both miRNA and mRNA data using ComBat-seq from the *sva* package. Using paired miRNA and mRNA data, two-tailed Spearman correlation tests were conducted for each putative miRNA–mRNA pair and corrected for false discovery rate (**FDR**) using the Benjamini-Hochberg procedure [47]. Based on biological evidence showing that miRNAs post-transcriptionally repress gene expression [48], we pruned the network to only retain miRNA–mRNA pairs that were negatively correlated at the FDR<0.05 level for downstream analyses (**Figure S2**). All network and subnetwork visualizations were generated using the *R* packages *igraph* (2.3.2) and *ggplot2* (4.0.3) [49,50] or with Cytoscape (3.10.4) software [51].

### 2.7. Differential miRNA Expression Analysis

The UMI count data were filtered to include only miRNA transcripts, normalized by computing logCPM, and filtered to remove miRNAs with low expression (logCPM<0). Differentially expressed miRNAs (**DEmiRs**) were identified using the limma-voom pipeline in the *edgeR* package [36,40,41]. This pipeline uses empirical Bayes moderated t-statistics to improve statistical power to detect differentially expressed transcripts while maintaining a low false discovery rate [52]. We included coefficients for confounding and precision variables including maternal race, maternal education, maternal smoking status, self-reported maternal alcohol consumption, household income (continuous), maternal pre-pregnancy body mass index (continuous), maternal age (continuous), labor status, sequencing batch, and fetal sex based on a directed acyclic graph (**DAG**, **Figure S3)**. Racism is a fundamental cause of health inequities in perinatal outcomes, including sPTB [53–55]. Maternal self-reported race was included in our models as a proxy of racism and structural inequities. We also conducted an analysis to identify DEmiR associated with gestational length (continuous, weeks) as a secondary outcome. We considered DEmiR significant with FDR<0.05.

We used miRbase (release 22.1) coordinates to identify primary miRNA transcripts located in placenta-specific miRNA loci, including the C14MC within the *DLK1*-*DIO3* locus within the 14q32 region and two clusters within the 19q13.42 region, C19MC [56,57] and miR-371 family [22,58].

### 2.8. Enrichment Analysis

We conducted enrichment analysis using the one-sided Fisher’s exact test to determine if each miRNA’s regulon, or all the genes regulated by a given miRNA, was enriched in specific Kyoto Encyclopedia of Genes and Genomes (**KEGG**) gene set pathways [59]. KEGG pathways (release 111.0) were limited a priori to exclude human disease or drug development pathways. We considered the target genes within our miRNA–mRNA network as the background list of genes for our over-representation analysis. This is because nonspecific background lists bias over-representation tests away from the null [60]. We considered pathways with FDR<0.05 to be significant.

To identify sPTB-enriched miRNA regulons, we used a previously published transcriptomic signature of sPTB generated from RNA-sequencing data in a joint analysis of the CANDLE and the Global Alliance to Prevent Prematurity and Stillbirth (**GAPPS**) cohorts [36]. We performed miRNA enrichment analyses on these sPTB signatures to identify miRNAs enriched for gene lists, in which we used each miRNA’s regulon as a gene group, analogous to a KEGG pathway. In that study, 961 protein-coding genes and long non-coding RNA were significantly associated with sPTB at FDR<0.05 [36]. We used the protein-coding target genes within our miRNA–mRNA network as background for this over-representation analysis. We considered miRNAs with FDR<0.05 to have regulons significantly enriched for sPTB genes.

### 2.9 *In vitro* validation of oxidative phosphorylation differences in selected miRNAs associated with sPTB

We performed miRNA inhibition experiments for selected miRNAs using the extravillous trophoblast HTR-8/SVneo placental cell line (ATCC, CRL-3271). Three C14MC miRNAs associated with sPTB (hsa-miR-376c-3p, hsa-miR-377-3p, and hsa-miR-381-3p) were selected for downstream analysis. Cells were maintained according to ATCC guidelines at 37°C with 5% CO2 and ambient O2 in 10-cm tissue culture dishes using RPMI-1640 media supplemented with 10% fetal bovine serum, 1% penicillin-streptomycin, 1 nM sodium pyruvate, and 10 nM HEPES.

miRNA inhibition experiments were performed in 96-well plates with 2,000 HTR-8/SVneo cells per well at passage 21. Separate transfections were performed using the ThermoFisher mirVana miRNA inhibitor assay for hsa-miR-376c-3p (Assay #MH12260), hsa-miR-377-3p (#MH10524), and hsa-miR-381-3p (#MH10242) or mirVana miRNA inhibitor negative control (#4464076). Inhibitors were reconstituted in nuclease-free water to a stock solution of 100 μM. Each inhibitor was diluted to a final concentration of 150 nM in nuclease-free water and transfected into cells using Lipofectamine RNAiMax reagent (ThermoFisher Scientific, #13778075) per manufacturer’s instructions. Cells were incubated for 6 hours with lipofectamine/miRNA inhibitor complex containing media before receiving a media change to fresh RPMI-1640 complete media for the remainder of the 48-hour incubation.

#### 2.9.1. Seahorse Oxidative Phosphorylation Assay

After completion of 48-hour transfection, oxygen consumption rate (**OCR**) parameters were determined using the Agilent Seahorse XF Cell Mito Stress Test Kit (Agilent Technologies, #103015-100) and a Seahorse XFe96 Analyzer (Agilent Technologies, Santa Clara, CA) according to manufacturer’s protocol. All assay media and oxygen consumption reagents were prepared on the same day as the assay. Prior to the assay, the 96-well plate was washed twice with XF RPMI assay media and allowed to equilibrate for 45-60 minutes in a 37°C cell culture oven. OCR data acquisition and normalization was completed using Wave Software (Agilent Technologies, v2.6.3) with the included XF Cell Mito Stress Test Kit assay template. Oligomycin (final concentration: 1.5 μM), FCCP (final concentration: 1.0 μM), and rotenone/antimycin A (0.5 μM) were sequentially added according to the experimental protocol. Upon completion of data acquisition, all wells were washed twice with 200 μL of 1X PBS, pH 7.4, and lysed in 200 μL of RIPA buffer. All lysed samples were stored in a pre-labeled 1.5 mL microcentrifuge tube at -80°C until completion of bicinchoninic acid (**BCA**) assay for protein quantitation.

A Pierce BCA Assay (ThermoFisher Scientific, #23227) was completed per the manufacturer’s instructions with all standards analyzed in triplicate and all unknowns analyzed in duplicate. BCA absorbance was measured at 562 nm and a standard curve was generated to calculate protein concentrations in ug/mL. To account for well-to-well variations, OCR parameters were normalized using protein concentrations.

All OCR parameters, including basal respiration, proton leak-linked respiration, ATP-linked respiration, maximal respiration, reserve capacity, and ATP coupling efficiency, were calculated using manufacturer recommendations as previously described [61]. Two sample t-tests were completed comparing each miRNA inhibitor group to the negative control (*n* = 4 per condition).

## 3. RESULTS

### 3.1. Descriptive Characteristics of Cohort

This analysis was conducted using paired mRNA and microRNA sequencing data (*N*=765) from placental samples collected within the CANDLE study [37]. To identify miRNAs associated with sPTB and gestational length, we excluded pregnancies with induced labor or indications for medically induced birth (**Figure S1**). Of 439 participants, 35 (8%) pregnancies resulted in sPTB (**Table 1**). The average age of participants was 28 years, 54.0% of participants self-identified as Black/African American, 40.1% identified as white, and there was no difference in fetal sexes.

Participants who delivered preterm had lower pre-pregnancy BMI and household income compared to participants that delivered at term, and no differences were observed by self-reported race, education, or age. Because sPTB follows preterm labor or preterm pre-labor rupture of membranes, participants were more likely to experience labor and deliver vaginally.

### 3.2. Generating a Placenta-specific miRNA–mRNA network

We linked miRNAs to their computationally predicted target genes and validated these associations with paired miRNA and mRNA sequencing data from CANDLE participants to construct the miRNA–mRNA network (**Figure S1**). Placentally expressed miRNAs (563 miRNAs) regulated 91.3% of placentally expressed protein-coding genes within our dataset. We used miRbase miRNA genome coordinates to identify miRNA located in placenta-specific miRNA clusters, including the C14MC (54 genes), C19MC (46 genes), and miR-371 family (4 genes). Our network has high coverage of these placenta-specific miRNA clusters, including miRNA from 46 C14MC genes, 45 C19MC genes, and 6 miR-371 family genes.

We have made the full results of the miRNA–mRNA network available in a searchable web application, Placenta miRNA–mRNA Regulatory Network, which can be accessed at https://www.paquettelab.org/placental-computational-tools/mirnanetwork. This application includes 4 tools: (i) a target gene search tool, (ii) a miRNA search tool, (iii) a KEGG pathway lookup tool, and (iv) an enrichment test tool. The search tools allow users to search the model for specific target genes or miRNAs of interest and evaluate individual relationships between miRNAs and target genes. The KEGG lookup tool allows users to look up KEGG Pathway gene sets that are enriched for members of a miRNA’s regulon by searching the results of our KEGG pathway enrichment analysis. The enrichment test tool allows users to import a list of genes of interest (such as the list of sPTB-associated genes presented here) and identify miRNAs whose target gene lists contain overrepresentation of the genes of interest. The entire miRNA–mRNA network is provided in **Table S2** and the results of the KEGG pathway enrichment in **Table S3**, for those who interested in accessing the network separate from the web application.

### 3.3. Shared and distinct miRNA signatures of sPTB and Gestational Length

We identified 19 differentially expressed miRNAs (DEmiRs) using covariate-adjusted linear regression models (**Figure 1**). 7 microRNAs were positively associated, and 12 were inversely associated with sPTB (**Figure S4**). These DEmiRs were from 14 microRNA families identified in TargetScan, including 4 microRNAs (hsa-miR-376a-3p, hsa-miR-376c-3p, hsa-miR-377-3p, and hsa-miR-381-3p) from the C14MC miRNA cluster, and 4 miRNAs from the let-7 family. 20 DEmiR associated with gestational length (FDR<0.05, **Table 2**), including 10 miRNAs that were associated with both gestational length and sPTB.

**Figure 1.**
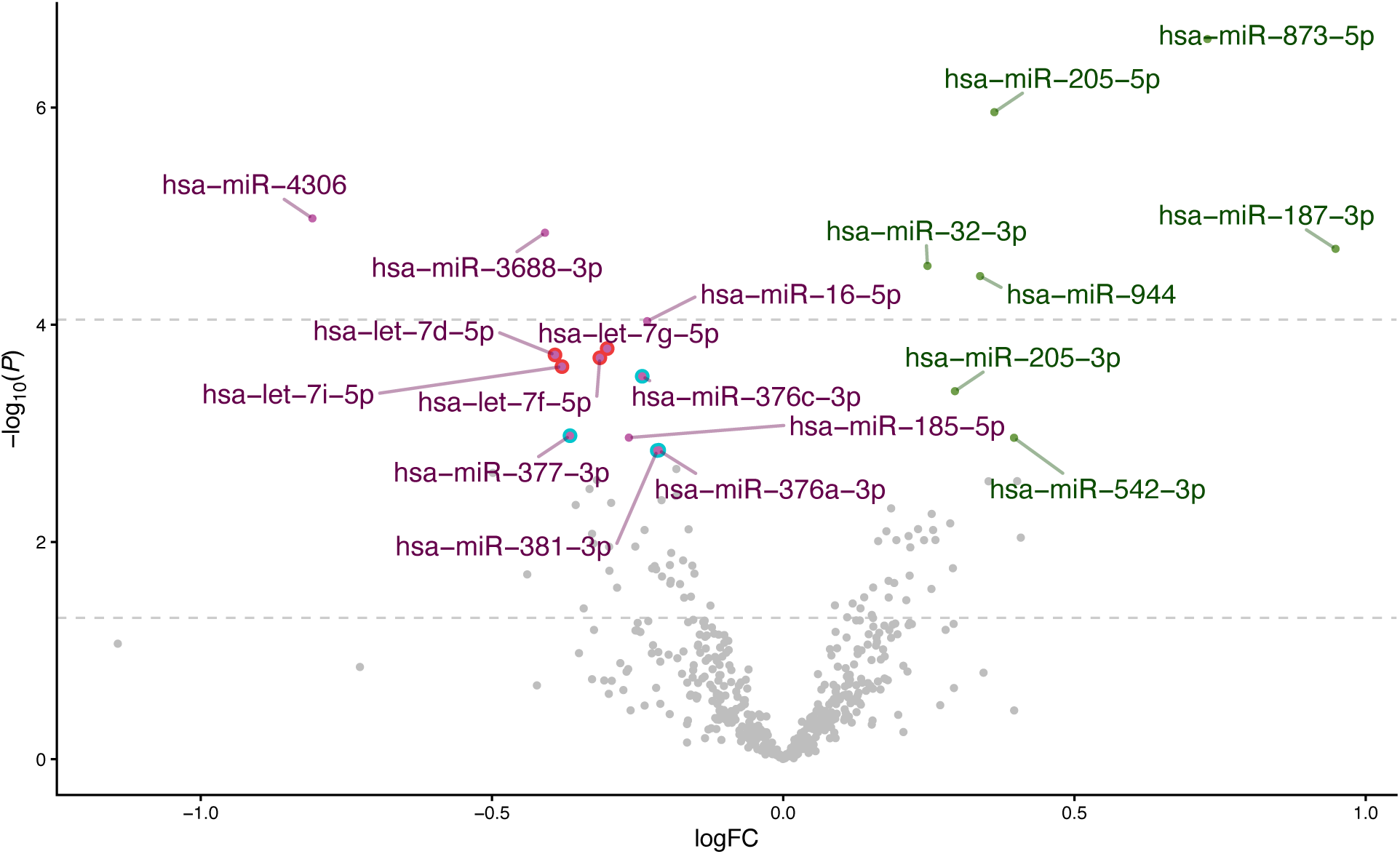
Volcano plot of spontaneous preterm birth (sPTB) associated differentially expressed microRNA (DEmiR). Members of the let-7 family are outlined in red, and members of the chromosome 14 microRNA cluster are outlined in blue. DEmiR was statistically defined by FDR<0.05. DEmiR with increased expression in sPTB are shaded green and those with decreased expression in sPTB are shaded magenta.

**Table 2.** Significant associations between miRNA and spontaneous preterm birth (sPTB) or gestational length in weeks.

| miRNA | sPTB |  | Gestational Length (weeks) |  | MiRNA Regulon Characteristics |  |
| --- | --- | --- | --- | --- | --- | --- |
|  | log<sub>2</sub> FC | FDR | log<sub>2</sub> FC | FDR | Regulon size | sPTB-associated targets<sup>a</sup> |
| hsa-let-7f-5p | -0.315 | 0.010 | 0.049 | 0.060 | 572 | 30 |
| hsa-let-7g-5p | -0.302 | 0.010 | 0.050 | 0.050 | 636 | 30 |
| hsa-let-7i-5p | -0.380 | 0.011 | 0.060 | 0.060 | 635 | 29 |
| hsa-miR-185-5p | -0.265 | 0.036 | 0.034 | 0.216 | 500 | 21 |
| hsa-miR-376a-3p | -0.214 | 0.042 | 0.032 | 0.152 | 103 | 6 |
| hsa-miR-376c-3p | -0.242 | 0.013 | 0.040 | 0.058 | 176 | 9 |
| hsa-miR-377-3p | -0.366 | 0.036 | 0.062 | 0.076 | 203 | 16 |
| hsa-miR-381-3p | -0.216 | 0.042 | 0.033 | 0.152 | 503 | 47 |
| hsa-miR-542-3p | 0.396 | 0.036 | -0.067 | 0.086 | 411 | 63<sup>b</sup> |
| hsa-let-7d-5p | -0.392 | 0.010 | 0.071 | 0.023 | 658 | 30 |
| hsa-miR-16-5p | -0.234 | 0.006 | 0.042 | 0.016 | 554 | 15 |
| hsa-miR-187-3p | 0.948 | 0.002 | -0.222 | 3.52 × 10<sup>-4</sup> | 32 | 1 |
| hsa-miR-205-3p | 0.295 | 0.016 | -0.060 | 0.016 | 926 | 79 |
| hsa-miR-205-5p | 0.362 | 3.07 × 10<sup>-4</sup> | -0.059 | 0.010 | 660 | 46 |
| hsa-miR-32-3p | 0.248 | 0.003 | -0.047 | 0.010 | 1317 | 46 |
| hsa-miR-3688-3p | -0.409 | 0.002 | 0.066 | 0.016 | 123 | 10 |
| hsa-miR-4306 | -0.808 | 0.002 | 0.134 | 0.016 | 533 | 23 |
| hsa-miR-873-5p | 0.728 | 1.31 × 10<sup>-4</sup> | -0.169 | 2.05 × 10<sup>-6</sup> | 16 | 0 |
| hsa-miR-944 | 0.338 | 0.003 | -0.055 | 0.025 | 860 | 48 |
| hsa-let-7a-5p | -0.184 | 0.059 | 0.042 | 0.016 | 33 | 1 |
| hsa-miR-122-5p | -0.328 | 0.123 | 0.087 | 0.018 | 98 | 8 |
| hsa-miR-126-3p | -0.163 | 0.120 | 0.044 | 0.016 | 3 | 0 |
| hsa-miR-139-5p | -0.254 | 0.130 | 0.074 | 0.016 | 587 | 77<sup>b</sup> |
| hsa-miR-147b-3p | 0.408 | 0.124 | -0.127 | 0.010 | 6 | 0 |
| hsa-miR-183-5p | 0.255 | 0.099 | -0.067 | 0.016 | 99 | 4 |
| hsa-miR-218-5p | 0.211 | 0.259 | -0.064 | 0.040 | 117 | 10 |
| hsa-miR-32-5p | 0.242 | 0.124 | -0.068 | 0.016 | 596 | 7 |
| hsa-miR-378a-5p | 0.352 | 0.064 | -0.087 | 0.016 | 1056 | 27 |
| hsa-miR-425-3p | 0.019 | 0.872 | -0.029 | 0.023 | 31 | 1 |

Differentially expressed miRNA were identified using linear models adjusted for maternal race, maternal education, maternal smoking status, maternal alcohol consumption, maternal income, maternal pre-pregnancy body mass index, maternal age, labor status, sequencing batch, and fetal sex. FDR<0.05 was considered significant. The table is arranged with the distinct signatures of sPTB, the shared signatures of sPTB and gestational length, and the distinct signatures of gestational length. FC, fold change; FDR, false discovery rate; sPTB, spontaneous preterm birth.
<sup>a</sup> sPTB-associated protein-coding genes were identified in a previous study using FDR<0.05 [36]
<sup>b</sup> The miRNA regulon is significantly enriched for sPTB target genes (Fisher's exact test, FDR<0.05)

### 3.4. Generating a miRNA–mRNA subnetwork of sPTB

We also identified which of the mRNA targets of DEmiRs were also associated with sPTB from our previously published placental transcriptomic signature of sPTB, (938 protein-coding genes) [36]. We used the miRNA–mRNA network that we constructed to link the 19 sPTB-associated DEmiR to 4,433 target genes, including 317 that were previously associated with sPTB (**Figure 2**). These sPTB-associated target genes included 22 TFs and 295 other protein-coding genes, representing 33.8% (317/938) of the sPTB protein-coding gene signature. The miR-542-3p regulon (411 genes) was significantly enriched for sPTB-associated genes (63/411, Fisher’s exact test, FDR<0.05). We integrated the 22 sPTB-associated TFs with our placental transcriptional regulatory network [62] to find that these TFs in turn regulate an additional 262 sPTB-associated target genes. Thus, the 18 sPTB-associated miRNA directly or indirectly regulate 61.7% of the sPTB signature (579/938 genes), including 22 sPTB-associated TFs and 557 sPTB-associated target genes.

**Figure 2.**
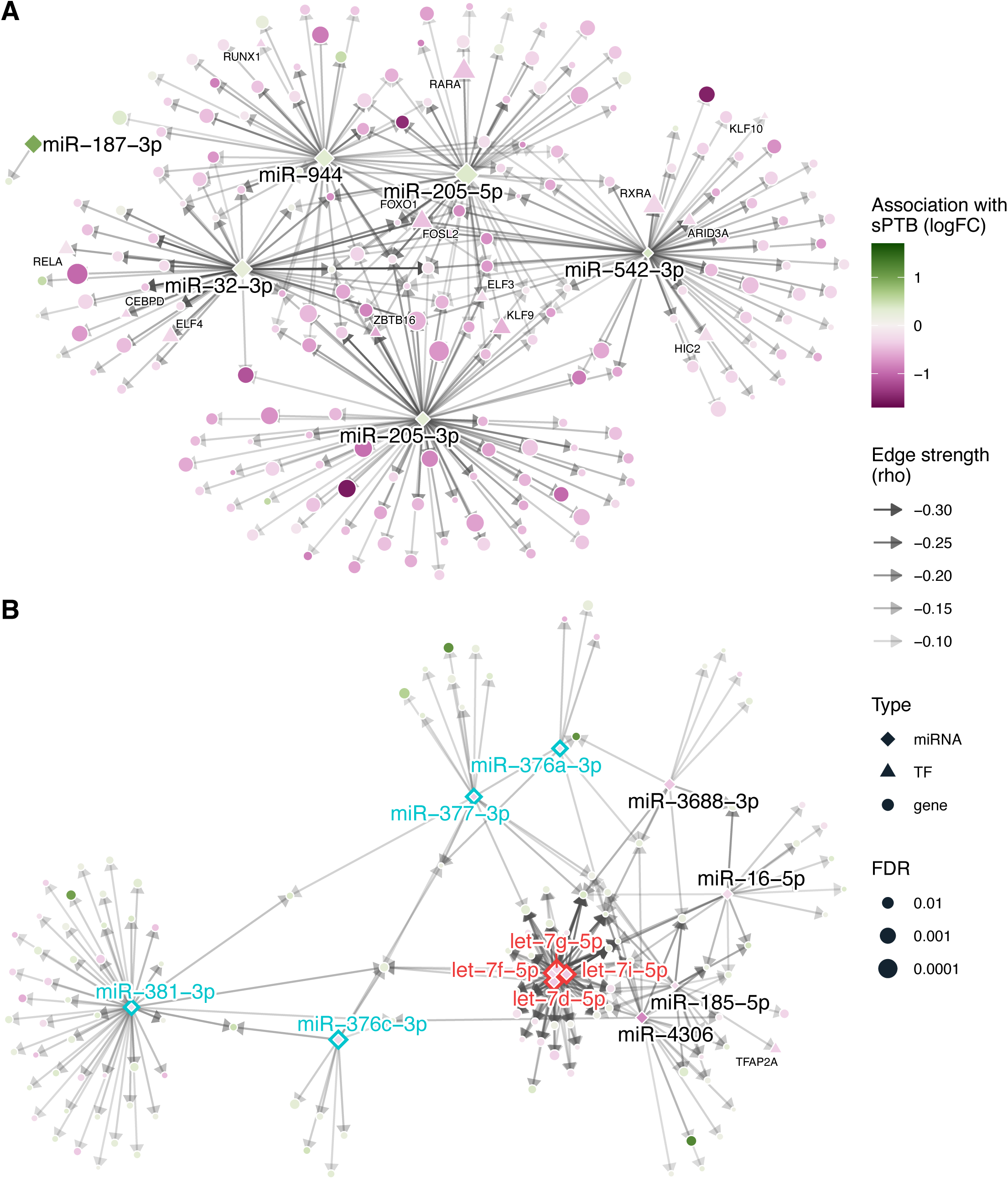
The subnetwork of sPTB-associated miRNA and their sPTB-associated target genes. The (**A**) up-regulated miRNA and (**B**) downregulated miRNA and their target genes were laid out according to a forced-directed algorithm based on edge strength (Spearman’s *ρ* between each miRNA and target gene). The miRNA (diamonds) regulate genes identified as transcription factors (TF, triangles) and other protein-coding genes (circles). miRNA and TF symbols are labeled. Node color corresponds to the effect size (logFC) of the association with sPTB and node size corresponds to the negative log_10_-transformed false discovery rate (FDR). Members of the chromosome 14 miRNA cluster are labeled in blue and members of the let-7 family are labeled in red. Edge transparency corresponds to the edge strength.

The 6 miRNAs that were up-regulated in sPTB targeted 203 sPTB-associated target genes which were largely down-regulated in sPTB (186 genes, **Figure 2A**). The target genes were enriched for the KEGG pathway *Th17 cell differentiation* (Fisher’s exact test, FDR<0.05) and 12 other pathways at the FDR<0.1 threshold (**Figure S5**), including immune, signal transduction, and developmental and regenerative pathways. The 12 miRNAs that were down-regulated in sPTB targeted 127 sPTB-associated genes and this set of target genes was not significantly enriched for any KEGG pathways (Fisher’s exact test, FDR>0.1).

The C14MC-derived hsa-miR-377-3p was downregulated in sPTB and was enriched for the KEGG pathways *oxidative phosphorylation* (9/99 genes) and *cardiac muscle contraction* (6/43 genes). The 4 sPTB-associated miRNA in the C14MC family together regulated 825 genes and were significantly enriched for genes involved in *oxidative phosphorylation* (21/99 genes) (FDR<0.05) (**Figure 3**). Five of the genes in the oxidative phosphorylation (**OXPHOS**) pathway were upregulated in sPTB in our previous study (*ATP6V1H, NDUFA10, NDUFA5, NDUFA6, PPA2*) [36]. The four microRNAs from the let-7 family associated with sPTB contained 764 genes but were not significantly enriched for any KEGG pathways (FDR>0.1). Given the unique expression pattern of the C14MC family in the placenta, we further investigated the subnetwork of C14MC and their predicted mRNA targets. The subnetwork includes 67 mature miRNAs from 46 genes, which together target 5460 protein-coding predicted target genes validated in our paired miRNA–mRNA analysis. In this subnetwork, 11 miRNA had regulons significantly enriched for 18 KEGG pathways related to protein synthesis, chromatin regulation, cellular metabolism, and cellular damage control (**Figure S6**).

**Figure 3.**
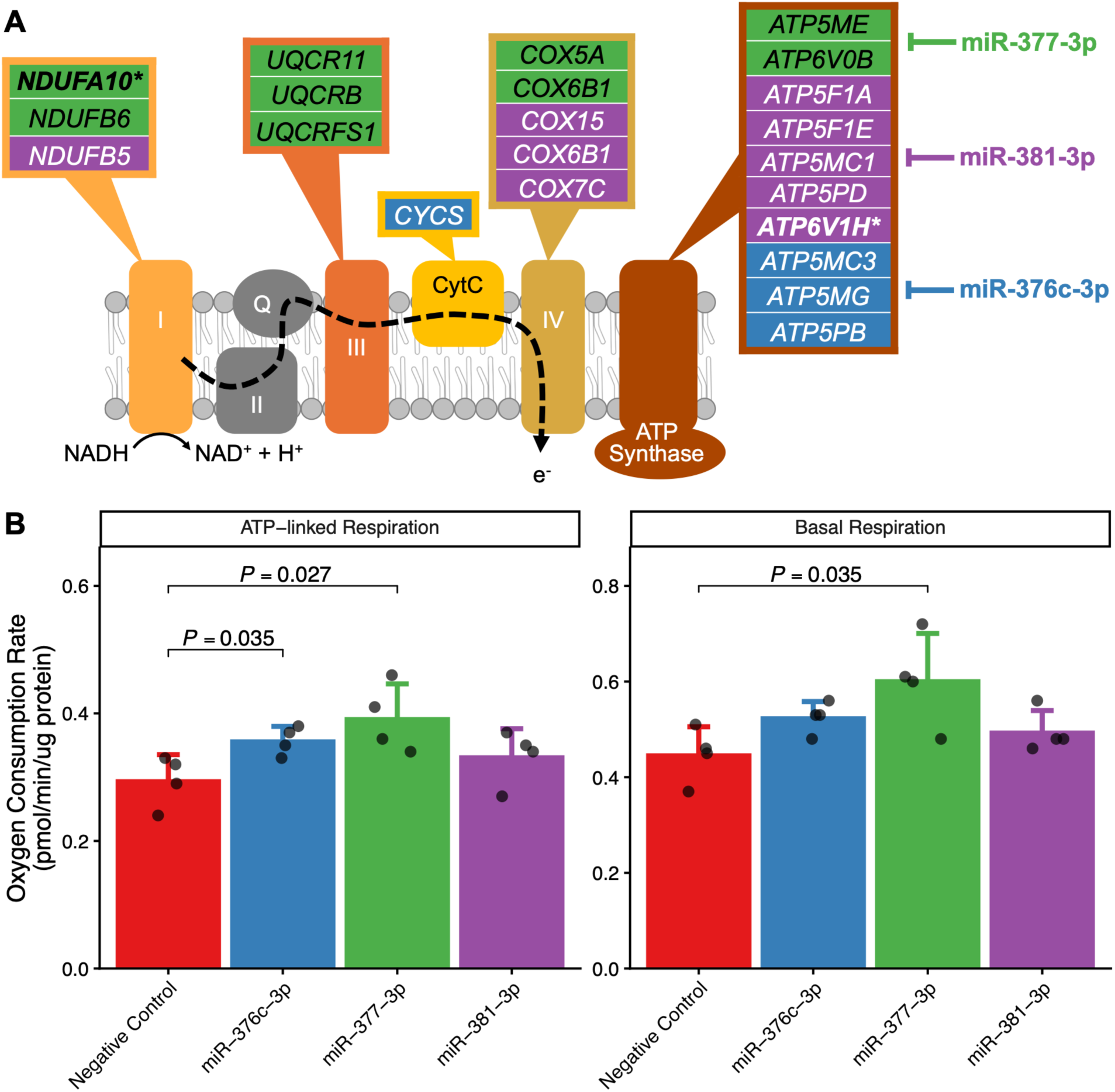
Mitochondrial respiration following microRNA inhibition in placental cells. (A) We identified target genes of miR-376c-3p, miR-377-3p, and miR-381-3p, which are colored blue, green, and purple, respectively. We previously reported on 5 genes in this pathway that were differentially expressed in sPTB and all 5 were significantly upregulated [36]. Two of these sPTB-associated genes were also regulated by C14MC miRNA (*NDUFA10, ATP6V1H*), and these are bolded and denoted by *. (B) Differences between oxygen consumption rates between the negative control and each microRNA inhibition were assessed independently using two-sample T-tests. Only comparisons with *P*<0.05 are labeled.

### 3.5. Validating OXPHOS as a pathway regulated by sPTB-associated C14MC

The miRNA–mRNA network revealed that sPTB-associated C14MC hsa-miR-376c-3p, hsa-miR-377-3p, and hsa-miR-381-3p targeted genes involved in the OXPHOS pathway (**Figure 4**). The targets identified in this analysis included subunits of electron transport chain Complexes I, III, and V (ATP synthase). Thus, we hypothesized that inhibiting any of these miRNAs would increase basal respiration and ATP-linked respiration measured using the Seahorse OXPHOS assay. Inhibition of hsa-miR-377-3p significantly increased the basal respiration rate (**Figure 3**, *P* = 0.035) and ATP-linked respiration rate (**Figure 3**, *P* = 0.027). Inhibition of hsa-miR-376c-3p significantly increased the ATP-linked respiration rate (**Figure 3**, *P* = 0 .035) and tended to increase the basal respiration rate, albeit at marginal significance (*P* = 0.059). We observed no other significant associations with OCR parameters of the OXPHOS pathway, including proton leak-linked respiration, maximal respiration, reserve capacity, and ATP coupling efficiency.

## 4. DISCUSSION

This comprehensive assessment used paired placental miRNA and mRNA sequencing in a large, well characterized pregnancy cohort to identify differences in the placental miRNA profiles related to sPTB. Our principal findings were: 1) 19 miRNAs associated with sPTB that directly regulate more than 33% of placental protein-coding genes associated with sPTB, 2) increased expression of miRNAs targeting sPTB-associated protein-coding genes involved in immune pathways, 3) decreased expression of genes in the placenta-specific C14MC family that were involved in oxidative phosphorylation. Our findings are consistent with a prior report of placental miRNAs in sPTB [34], and expand on them with well-defined spontaneous phenotypes, adjustment for confounding variables, and leveraging paired miRNA and mRNA expression data to better characterize miRNA regulation in the placenta. Taken together, these findings yield novel insight into the molecular regulation of sPTB.

Our sPTB microRNA–mRNA subnetwork revealed that sPTB microRNA directly or indirectly targeted 61.7% of differentially expressed genes associated with sPTB. This highlights a relatively small number of miRNAs as potential regulators of sPTB, which can inform future work into promising targets and biomarkers. Our results are consistent with another recent analysis of miRNA expression in the basal plate of the placenta in sPTB (*n*=6) compared to spontaneous term births (*n*=6) [34]. That study reported that hsa-miR-187-3p was positively associated with sPTB and hsa-miR-376a-3p, hsa-miR-376c-3p, and hsa-miR-377-3p were inversely associated with sPTB [34], all of which were replicated in the present analysis.

We established a placenta-specific miRNA–mRNA network to understand the biological pathways the sPTB-associated miRNAs participated in. Six miRNAs with higher expression in sPTB placentas, including hsa-miR-542-3p, regulated of 21.6% of the sPTB-associated target genes, which were enriched for the *Th17 cell differentiation* pathway and other immune and cell signaling pathways involved in cell-mediated immunity. These pathways were down-regulated at the gene expression level, which is counterintuitive because Th17 cells participate in inflammatory responses and have been implicated in preterm labor [63]. However, immune system activation also triggers anti-inflammatory gene programs to enable resolution of the inflammatory [64]. In trophoblast cells, elevated hsa-miR-542-3p expression might play a role in inhibiting macrophage recruitment [65]. Thus, placental miRNA might play a role in regulating immune responses linked to sPTB pathogenesis.

Notably, we observed that hsa-miR-542-3p expression was higher in sPTB and its regulon was significantly enriched for sPTB-associated target genes. Hsa-miR-542-3p is a highly conserved mammalian miRNA that is necessary for implantation as well as inflammation [66,67]. It suppresses decidualization in endometrial stromal cells and inhibits angiogenesis [68,69]. Among the sPTB-associated targets of hsa-miR-542-3p were important placental transcription factors, including *ARID3A*, *FOSL2*, *FOXO1*, and *RXRA*. *ARID3A* is required for placental development, and participates in shaping placental structure, metabolism, and immune tolerance [70,71]. *FOSL2* is a member of the Fos family and serves as a subunit in the AP-1 transcription factor, which governs cell processes including differentiation, proliferation, and apoptosis [72]. *FOXO1* is an essential transcription factor directing placental morphology and decidualization [73,74]. *RXRA* is a transcription factor that forms heterodimers with many nuclear hormone receptors, with widespread impacts on gene expression [75]. Additionally, hsa-miR-542-3p was part of the distinct signature of sPTB, which was not also associated with gestational length. Taken together, this suggests that hsa-miR-542-3p might participate in an important regulatory circuit involved in sPTB pathogenesis, potentially through placental implantation or development.

The C14MC is a paternally imprinted, maternally expressed miRNA cluster that is highly conserved across placental mammals and is located within the *DLK1*-*DIO3* locus in humans [76]. We report that 4 C14MC-derived miRNAs (hsa-miR-376a-3p, hsa-miR-376c-3p, hsa-miR-377-3p, and hsa-miR-381-3p) were downregulated in sPTB in the villous placenta. Our results are consistent with another recent analysis demonstrating a pattern of lower expression from C14MC-derived miRNA in the basal plate of the placenta in sPTB [34]. These placental microRNAs have also been associated with other pregnancy pathologies including preeclampsia and placental accreta [77–79]. The roles for many of the C14MC-derived miRNA in placental development remain to be elucidated [76], though hsa-miR-376c-3p and hsa-miR-377-3p might regulate placental invasion and growth, respectively [77–80].

We identified OXPHOS gene targets for three C14MC members (hsa-miR-376c-3p, hsa-miR-377-3p, and hsa-miR-381-3p). In OXPHOS, cells oxidize nutrients sequentially through the complexes of the electron transport chain to produce ATP. Complexes I, III, and IV establish a proton gradient needed to drive ATP production at ATP synthase. Notably, Complexes I and III are sites of reactive oxygen species (**ROS**) generation [81]. Complex I also plays an important role as a regulator of the cellular NAD^+^/NADH ratio in the cell by oxidizing NADH to NAD^+^ during OXPHOS [81]. Hsa-miR-377-3p targeted subunits of Complex I (*NDUFA10, NDUFB6*), Complex III (*UQCR11, UQCRB, UQCRFS1*), Complex IV (*COX5A*, *COX6B1*), and ATP synthase (*ATP5ME, ATP6V0B*). Hsa-miR-376c-3p also targeted the electron carrier cytochrome C (*CYCS*) and ATP synthase subunits (*ATP5MC3*, *ATP5MG*, *ATP5PB*). Our functional validation results demonstrate that both hsa-miR-376c-3p and hsa-miR-377-3p inhibition increase the ATP-linked respiration rate and hsa-miR-377-3p inhibition also increases the basal respiration rate, consistent with our hypotheses. In normal pregnancy, placental energy metabolism favors glycolysis, even in aerobic conditions [82]. While aerobic glycolysis is less efficient than OXPHOS for ATP production, glycolytic metabolism generates energy, reducing equivalents, and precursors for macromolecule biosynthesis without producing ROS [83]. In contrast, we report lower expression of C14MC miRNAs targeting OXPHOS genes in sPTB, suggesting that OXPHOS might be upregulated in sPTB. In turn, ROS produced through OXPHOS could contribute to oxidative stress, which has been observed in sPTB and may serve as an underlying biological mechanism [84].

In humans, the let-7 miRNA family includes 10 mature miRNAs originating from the 5p-arm and we identified 5 that were differentially expressed in sPTB or with increasing gestational length. The let-7 miRNA family is also highly conserved among animals and regulates growth and differentiation in development [85]. Within the placenta, the let-7 family members are upregulated in the third trimester compared with the first trimester and are important regulators of growth [80,86]. In the JEG-3 and BeWo choriocarinoma cell lines, a let-7d inhibitor promoted proliferation, migration, and invasion [87]. Hsa-let-7d-5p is upregulated in placentas affected by preeclampsia compared to healthy controls [87]. The placenta has been shown to export miRNAs into maternal circulation, so monitoring circulating maternal miRNAs has been explored as a promising predictive test [21]. A case–control study of sPTB ≤ 32 weeks’ gestation compared to spontaneous term births identified and validated that expression of hsa-let-7g was upregulated in maternal plasma collected between 16 weeks and 19^+5^ weeks in sPTB cases [88]. Taken together with our findings in the C14MC, the pattern of downregulation among highly conserved miRNAs in sPTB compared to term placenta could reflect impaired placental function or dysregulated placental maturation leading to sPTB. Moreover, these miRNAs might serve as useful biomarkers to monitor in maternal plasma.

This work has several strengths. First, this research was conducted in a large, diverse, well-characterized pregnancy cohort, and rigorously accounted for a variety of confounding variables and multiple testing. Second, it leveraged paired mRNA and miRNA expression data within this cohort to generate a prototype placental miRNA–mRNA regulatory network in samples collected at delivery. We have made the full network available in a web application to enable other placental biology researchers to easily access this placenta-specific miRNA–mRNA network and use this database to identify miRNA regulators of their own gene lists of interest. Future directions for this research include external validation of the network with other cohorts.

This work should also be interpreted in the context of its limitations. First, despite rigorous adjustment for confounding variables, we cannot rule out the potential of residual confounding within this observational cohort. One important consideration is the role of prematurity in these findings, as sPTB is necessarily confounded by gestational length. Notably, we identified miRNA signatures of sPTB that were distinct from those associated with gestational length. Second, these data were generated in bulk tissue and cannot provide insight into cell-specific differences in miRNA expression in sPTB or in miRNA regulation. While large cohorts are not amenable to single-cell sequencing, future research should consider cellular deconvolution to capture cell-type specific differences [89]. Third, cross-sectional measurement of miRNA and mRNA might not fully capture temporal relationships between miRNA expression and mRNA degradation, which takes approximately 60 minutes from miRNA synthesis to mRNA target decay [90]. miRNA can also repress protein translation by binding mRNA, reducing protein abundance without accompanying changes in the transcriptome [90]. Finally, these findings are from a single cohort of participants predominantly self-identifying as Black, a population that is historically underrepresented in the scientific literature. In the future, these findings should be validated in additional cohorts.

In summary, this study identifies placental miRNAs that are differentially expressed in sPTB and with increasing gestational length. Notably, highly conserved miRNAs in the C14MC family are downregulated in sPTB, distinct from gestational length. Our epidemiological and in vitro validation suggest that hsa-miR-376c-3p and hsa-miR-377-3p might play a role in regulating oxidative phosphorylation. We also found that miRNAs targeted sPTB-associated protein-coding genes involved in immune pathways that have been implicated in the etiology of sPTB. Functional validation studies are needed to better understand the role of these miRNAs in regulating the placental transcriptome and the role of these miRNAs in placental function. Moreover, this study further provides insight into the regulatory role of placental miRNAs, which target a substantial portion of the placental genes associated with sPTB. Future work exploring these miRNAs as promising biomarkers of sPTB in blood is warranted.

## DECLARATIONS

### Ethics approval and consent to participate

All research activities for the Conditions Affecting Neurocognitive Development and Learning in Early childhood (CANDLE) Study were approved by the University of Tennessee Health Sciences Center IRB, the ECHO prenatal and early childhood pathways to health consortium (ECHO-PATHWAYS) single IRB and Seattle Children’s Research Institute (STUDY00004005).

### Availability of data and materials

Placental mRNA and miRNA sequencing data from CANDLE are available on dbGaP at accession number: phs003619.v1.p1. Covariate data are available by request from CANDLE. The full results of the miRNA–mRNA network available in web application, Placenta miRNA–mRNA Regulatory Network, which can be accessed at https://www.paquettelab.org/placental-computational-tools/mirnanetwork.

### Authors’ contributions

MP and AGP conceptualized the analysis. JM and TB curated the sequencing data. AGP, SS, KZL, NRB, and QZ secured funding. AGP, SS, and QZ provided supervision. MP constructed the network, conducted formal analysis, and visualized the data. EJF conducted in vitro experiments and created the web application. MP and AGP wrote the original draft. All authors contributed to reviewing and editing the final version. All authors read and approved the final manuscript.

## Acknowledgements

The authors would like to thank the study staff, laboratory technicians, data teams and co-investigators involved in the CANDLE and the ECHO-PATHWAYS consortium for their invaluable contributions. This manuscript has been reviewed by PATHWAYS for scientific content and consistency of data interpretation with previous PATHWAYS publications. The authors are also grateful to Timothy J. Cherry, PhD, for his guidance in the in vitro microRNA inhibition experiments.

## SUPPLEMENTAL INFORMATION

**Supplement 1** (PDF) contains Table S1 and Figures S1-S6. **Supplement 2** (XLSX) contains Table S2 and S3.

