## Supplement 1 for "Placental microRNA signatures of spontaneous preterm birth"

**Table S1**. Comparison of spontaneous births included in this analysis to the full CANDLE cohort.

| | Full CANDLE Cohort (*N*=1503) | Spontaneous Births in this analysis (*N*=439) | *P* value |
|---|---|---|---|
| **Maternal age at birth** | | | < 0.001 |
| Mean (SD) | 26.3 (5.4) | 27.8 (5.3) | |
| Missing | 75 | 0 | |
| **Maternal pre-pregnancy BMI** | | | 0.723 |
| Mean (SD) | 27.5 (7.5) | 27.4 (7.3) | |
| Missing | 5 | 0 | |
| **Maternal education** | | | < 0.001 |
| Less than high school | 184 (12.3%) | 31 (7.1%) | |
| High school graduate/GED | 709 (47.2%) | 191 (43.5%) | |
| Technical school | 138 (9.2%) | 38 (8.7%) | |
| College degree | 299 (19.9%) | 114 (26.0%) | |
| Grad/Professional degree | 171 (11.4%) | 65 (14.8%) | |
| Missing | 2 | 0 | |
| **Estimated total annual household income at enrollment** | | | < 0.001 |
| $0-4999 | 191 (14.0%) | 49 (11.2%) | |
| $5000-9999 | 105 (7.7%) | 36 (8.2%) | |
| $10000-14999 | 97 (7.1%) | 24 (5.5%) | |
| $15000-19999 | 100 (7.3%) | 25 (5.7%) | |
| $20000-24999 | 106 (7.7%) | 23 (5.2%) | |
| $25000-34999 | 155 (11.3%) | 50 (11.4%) | |
| $35000-44999 | 109 (8.0%) | 36 (8.2%) | |
| $45000-54999 | 106 (7.7%) | 45 (10.3%) | |
| $55000-64999 | 80 (5.8%) | 20 (4.6%) | |
| $65000-74999 | 85 (6.2%) | 42 (9.6%) | |
| $75000 or over | 234 (17.1%) | 89 (20.3%) | |
| Missing | 135 | 0 | |
| **Maternal self-reported race** | | | < 0.001 |
| Black/African American | 936 (62.4%) | 237 (54.0%) | |
| White | 467 (31.1%) | 176 (40.1%) | |
| Asian | 13 (0.9%) | 5 (1.1%) | |
| American Indian/Alaska Native | <5 | <5 | |
| Native Hawaiian/Other Pacific Islander | <5 | <5 | |
| Other | 6 (0.4%) | <5 | |
| Multiple race | 77 (5.1%) | 20 (4.6%) | |
| Missing | 2 | 0 | |
| **Maternal smoking status** [a] | | | 0.038 |
| Never | 1302 (86.7%) | 393 (89.5%) | |
| Previous or current smoker | 200 (13.3%) | 46 (10.5%) | |
| Missing | 1 | 0 | |
| **Maternal self-reported alcohol use** | | | 0.008 |
| No | 1381 (91.9%) | 391 (89.1%) | |
| Yes | 121 (8.1%) | 48 (10.9%) | |
| Missing | 1 | 0 | |

| | Full CANDLE Cohort (*N*=1503) | Spontaneous Births in this analysis (*N*=439) | *P* value |
|---|---|---|---|
| **Fetal sex** | | | 0.909 |
| Male | 736 (50.3%) | 220 (50.1%) | |
| Female | 726 (49.7%) | 219 (49.9%) | |
| Missing | 41 | 0 | |
| **Labor type** | | | < 0.001 |
| Spontaneous | 340 (23.4%) | 127 (28.9%) | |
| Spontaneous, augmented | 418 (28.7%) | 195 (44.4%) | |
| Induced | 448 (30.8%) | 0 (0.0%) | |
| No labor | 249 (17.1%) | 117 (26.7%) | |
| Missing | 48 | 0 | |
| **Delivery method** | | | 0.042 |
| Vaginal | 916 (62.9%) | 259 (59.0%) | |
| C-section | 540 (37.1%) | 180 (41.0%) | |
| Missing | 47 | 0 | |
| **Preterm birth** | | | 0.312 |
| Term | 1323 (90.9%) | 404 (92.0%) | |
| Preterm | 133 (9.1%) | 35 (8.0%) | |
| Missing | 47 | 0 | |

Continuous variables are reported as mean and standard deviation (SD) and differences were tested using two-sample T-tests. Categorical variables are reported as N (%) and were tested using $\chi^2$ tests.

[a] Maternal smoking status was a composite variable determined by self-report and urinary cotinine where available. Participants who self-reported ever smoking or who had urinary cotinine > 200 ng/mL were classified as previously or currently smokers

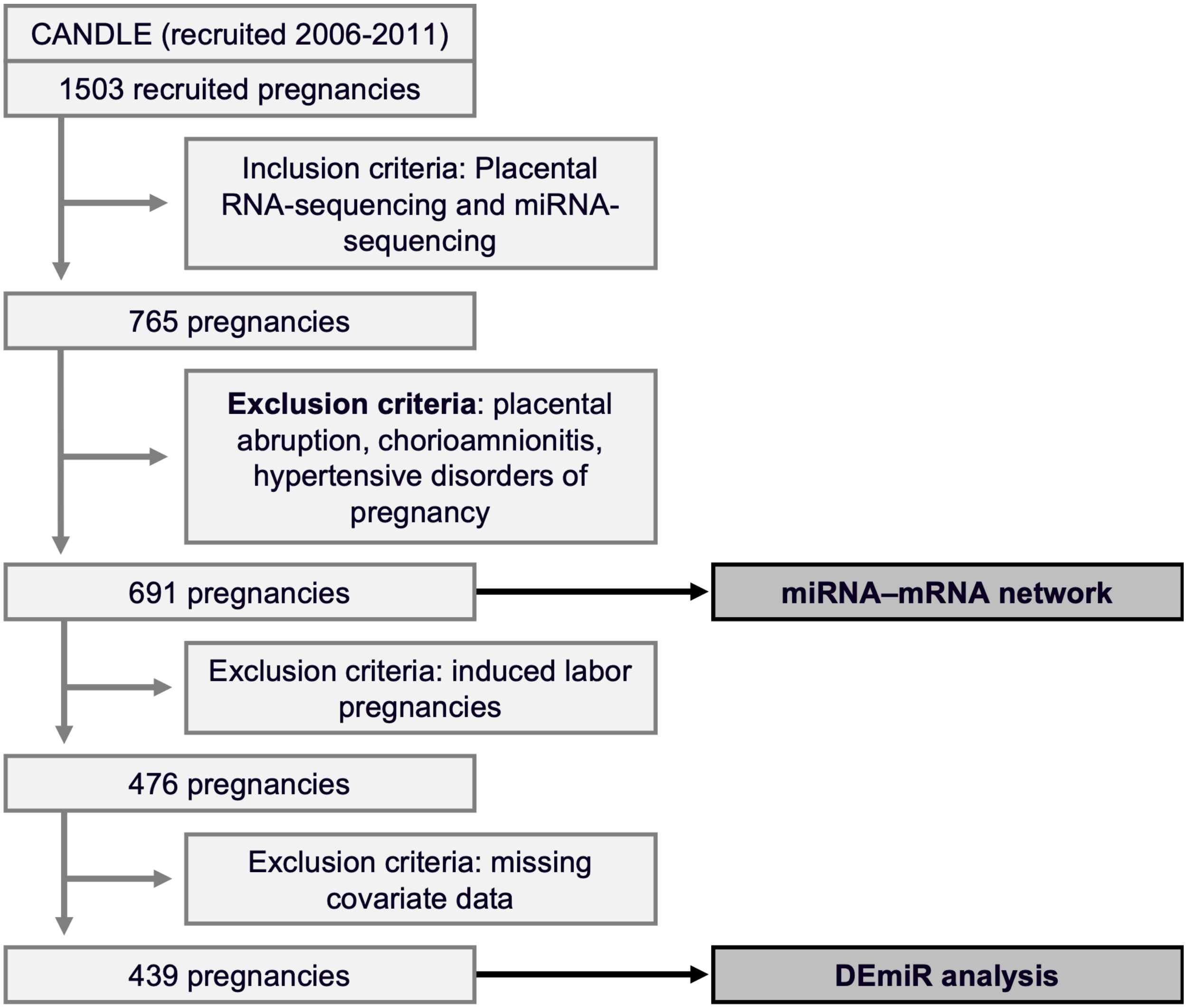


**Figure S1. CONSORT-style diagram of samples included in the analyses presented here**.

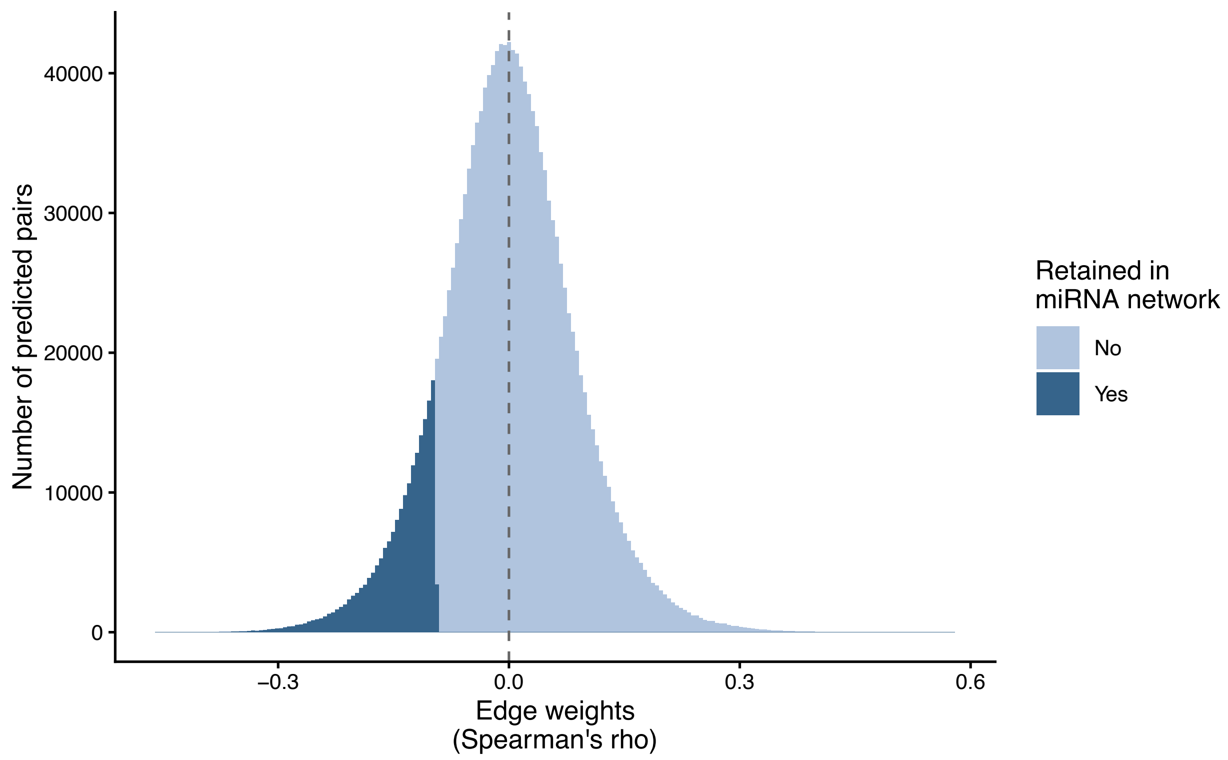


**Figure S2. Distribution of edge weights for computationally predicted miRNA-mRNA target pairs.** Edge weights for 1,586,889 computationally predicted miRNA–mRNA target pairs were tested using Spearman's rank correlation coefficient, rho, and corrected for false discovery rate (FDR). With a threshold of FDR<0.05, there were 200,168 significant positive correlations between predicted pairs and 199,118 significant negative correlations between predicted pairs. Since miRNA act by degrading target mRNA, we retained only the significant negative correlations for further analysis.

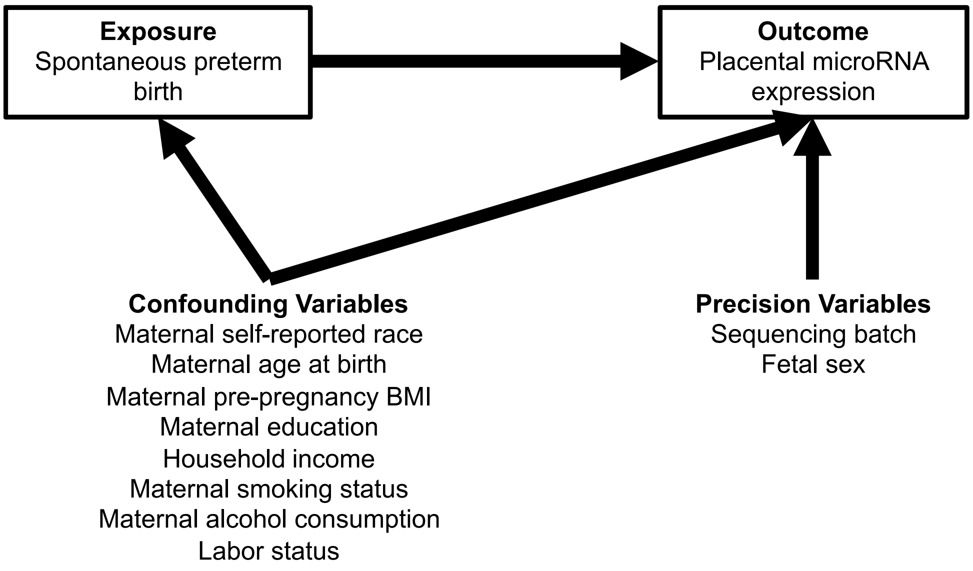


**Figure S3. Directed acyclic graph depicting covariates used in the analysis.**

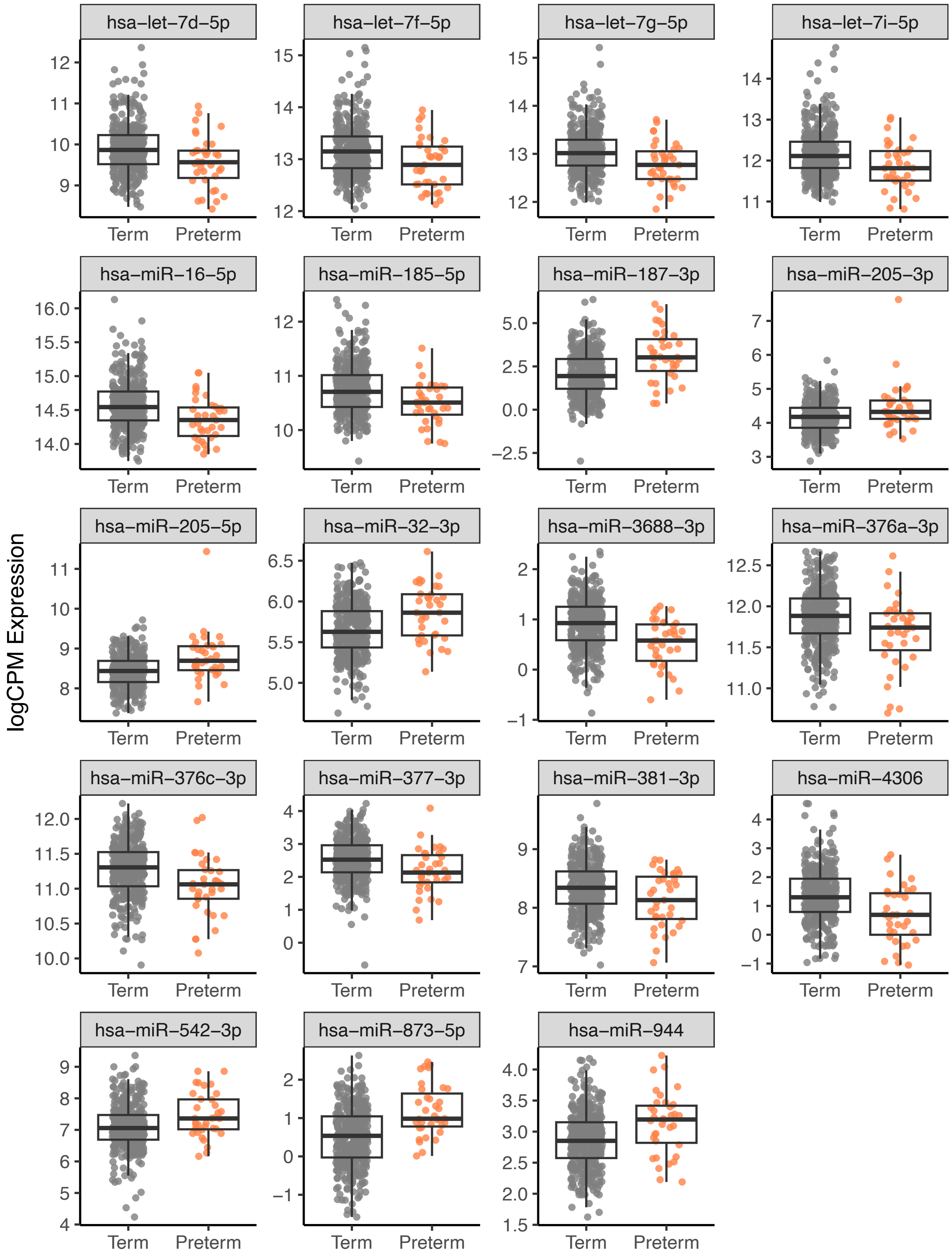


**Figure S4. Differential expression of miRNA in spontaneous preterm birth.** MiRNA expression is presented as logCPM for differentially expressed miRNA associated with spontaneous preterm birth (FDR<0.05). Each sample is plotted and boxplots represent the median and IQR for term and preterm samples.

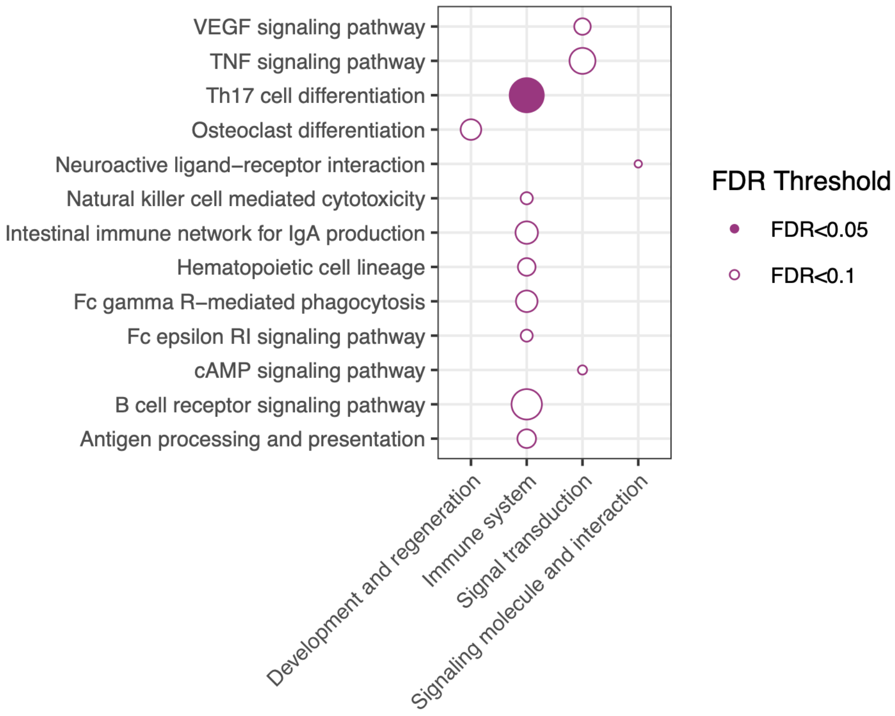


**Figure S5. MicroRNA upregulated in sPTB regulate target genes in pathways related to immune function.** The sPTB-associated target genes of the up-regulated miRNA were tested for KEGG pathway enrichment and corrected for false discovery rate (FDR). These sPTB-associated target genes were generally down-regulated in sPTB. Pathways are categorized according to KEGG pathway subheadings. All pathways were enriched (Fisher's exact test) at the FDR<0.10 level (open circles) or at the FDR<0.05 level (closed circles).

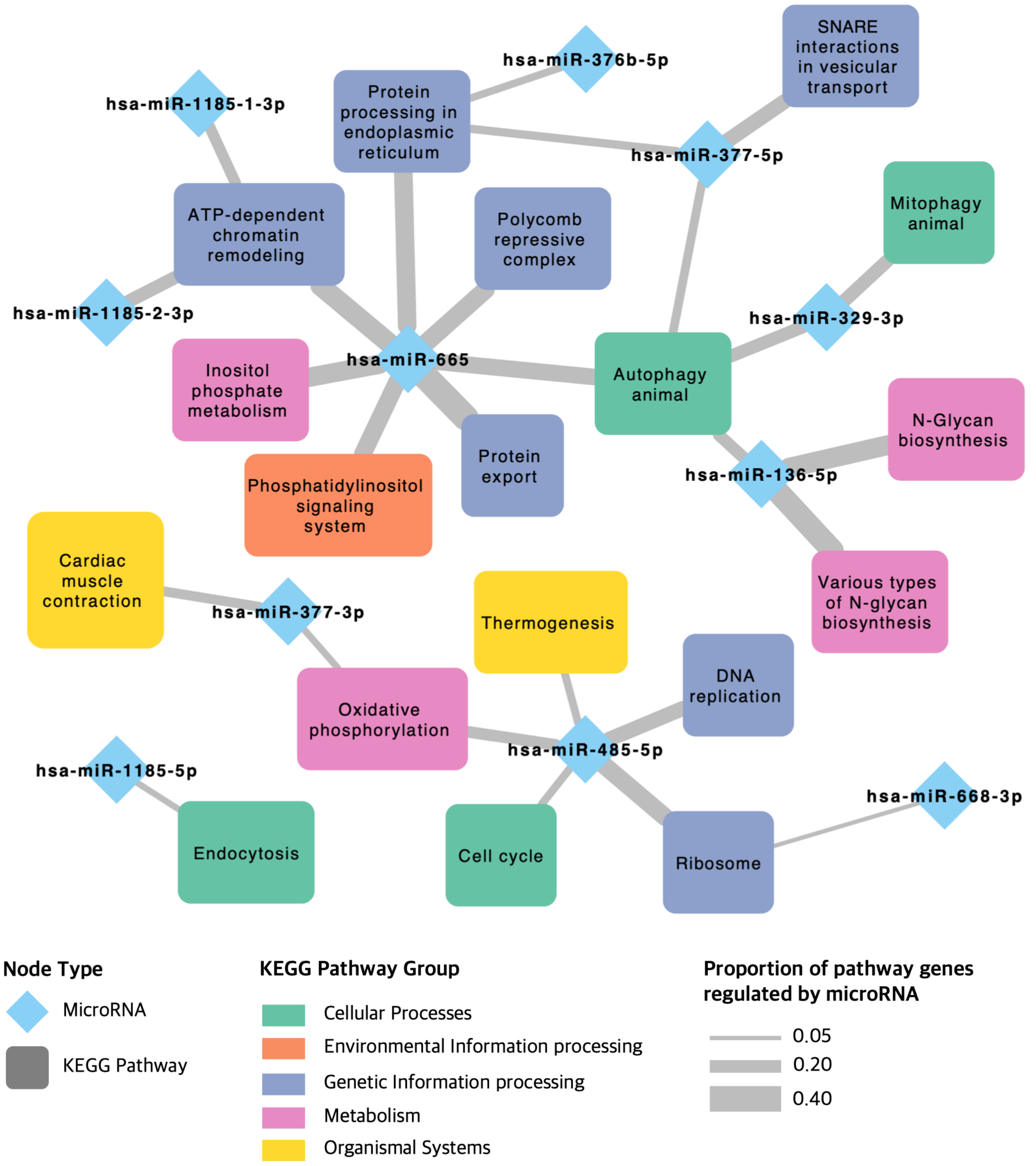


**Figure S6. C14MC microRNAs are enriched for KEGG pathways**. mRNA target genes for each C14MC member were tested for enrichment using KEGG pathways. Edges represent significant enrichment of target genes (FDR<0.05) and edge width corresponds to the proportion of pathway genes regulated by each C14MC member. Figure made in Cytoscape.
